# Microbiome Metabolome Integration Platform (MMIP): a web-based platform for microbiome and metabolome data integration and feature identification

**DOI:** 10.1101/2023.04.04.535534

**Authors:** Anupam Gautam, Debaleena Bhowmik, Sayantani Basu, Wenhuan Zeng, Abhishake Lahiri, Daniel H. Huson, Sandip Paul

**Affiliations:** Algorithms in Bioinformatics, Institute for Bioinformatics and Medical Informatics, University of Tübingen, Tübingen, Germany; International Max Planck Research School “From Molecules to Organisms,” Max Planck Institute for Biology Tübingen, Tübingen, Germany; Cluster of Excellence: EXC 2124: Controlling Microbes to Fight Infection, University of Tübingen, Tübingen, Germany; Cell Biology and Physiology Division, CSIR-Indian Institute of Chemical Biology, Kolkata, India; Academy of Scientific and Innovative Research (AcSIR), Ghaziabad-201002, India; Department of Computer Science, University of Illinois at Urbana-Champaign, Urbana, IL 61801, United States; Cluster of Excellence: EXC 2064: Machine Learning: New Perspectives for Science, University of Tübingen, Tübingen, Germany; Infectious Diseases and Immunology Division, CSIR-Indian Institute of Chemical Biology, Kolkata, India; Centre for Health Science and Technology, JIS Institute of Advanced Studies and Research Kolkata, JIS University, West Bengal, India

## Abstract

A microbial community maintains its ecological dynamics via metabolite crosstalk. Hence knowledge of the metabolome, alongside its populace, would help us understand the functionality of a community and also predict how it will change in atypical conditions. Methods that employ low-cost metagenomic sequencing data can predict the metabolic potential of a community, that is, its ability to produce or utilize specific metabolites. These, in turn, can potentially serve as markers of biochemical pathways that are associated with different communities. We developed MMIP (Microbiome Metabolome Integration Platform), a web-based analytical and predictive tool that can be used to compare the taxonomic content, diversity variation and the metabolic potential between two sets of microbial communities from targeted amplicon sequencing data. MMIP is capable of highlighting statistically significant taxonomic, enzymatic and metabolic attributes as well as learning-based features associated with one group in comparison with another. Further MMIP can predict linkages among species or groups of microbes in the community, specific enzyme profiles, compounds or metabolites associated with such a group of organisms. With MMIP, we aim to provide a user-friendly, online web-server for performing key microbiome-associated analyses of targeted amplicon sequencing data, predicting metabolite signature, and using learning-based linkage analysis, without the need for initial metabolomic analysis, and thereby helping in hypothesis generation.

## Introduction

Microorganisms are omnipresent and are important players in the global bio-geochemical cycles. They are also extensive symbionts for all major clades of life (1–4). The study of collective genetic material of microorganisms in any biome is the focus of microbiome research. This varied and intensifying research field is exploring the holistic influence of the microbial community in a diverse set of environments including ocean, soil, human body etc. In case of human health, the critical role of the microbiome has been fully recognized (5, 6). The microbial community structure and function is found to be governed by environmental factors, host physiology and interactions among the microbes (7–9). For a long time, the primary focus of microbiome studies has been to explore the differences in diversities across various sample groups (10). However, the current focus of microbiome research lies in gaining mechanistic insights into the functional capabilities of the microbiome and their impact on host physiology. The delicate balance of host-bacteria crosstalk depends on various factors, including microbial-origin molecules with hormone-like features, bacteria secreted factors like specific peptides and effector proteins, as well as host secreted factors such as IgA and antimicrobial peptides (11, 12). To understand the complex interactions between the microbial community and host physiology, the integration of metabolomics and microbiome analysis has emerged as a powerful approach. By examining the production, modification, or degradation of bioactive metabolites, this integrated analysis explores the intricate role of microbial communities in shaping host physiology. Disease-specific studies have utilized microbiome-metabolome integrative analysis followed by functional characterization to identify disease-associated microbial compounds (13–17). Integrating microbiome and metabolome data is crucial for researchers to unravel the complex web of interactions, identify disease-associated biomarkers, and gain a deeper understanding of the underlying mechanisms involved.

Several computational tools have emerged that attempt to unite the microbiome and metabolome data to either predict the metabolomic yield of a given microbial community or to study the intricate microbe-microbe or host-microbe interactions (18–22). MIMOSA, which is based upon Predicted Relative Metabolic Turnover (PRMT), connects metabolite levels and predicted microbial metabolic capabilities in metabolomes and metagenomes (18, 19). The application of machine learning models to predict metabolite composition from sequencing data was employed in MelonnPan (21). AMON aids in annotating metabolomics data by deciphering the source of the metabolites, either from microbes or from host (22). Here we introduce MMIP (Microbiome-Metabolome Integration Platform), an online platform designed to integrate and analyze microbiome and metabolome data. It is aimed at delineating differential community-level information and the metabolic potential of various microbial communities from targeted amplicon sequencing data utilizing the algorithms introduced in PRMT and MIMOSA. MMIP can emphasize statistically significant taxonomic, potential enzymatic and metabolic traits, as well as learning-based features associated with one group in comparison with another. Furthermore, we attempt to predict connections among the three layers of data (taxonomy abundance, predicted enzyme profiles and metabolite potential score) that discriminate between two microbial communities and possess probable biological significance.

## Materials and Methods

The MMIP web interface utilizes recent versions of bootstrap (v4.3.1) and HTML for a user-friendly experience. JavaScript (ECMA Script 2021) enables interactive plots and downloadable images. MMIP is composed of two distinct modules. Module-I is dedicated to comprehensive metagenomic analyses, encompassing taxonomic profile, diversity assessments, predicted enzyme profiles, metabolite potential evaluations, and feature identification. This module is adept at comparing these metagenomic analyses between two user-defined groups. Module-II, on the other hand, focuses on founding correlations between predicted metabolites with user-generated metabolomic data. It utilises the complete functionality of Module-I to generate the data necessary for conducting correlation analysis. Therefore, initiating Module-II will inherently yield correlation data between predicted metabolites with user-generated metabolomic data as well as the results derived from Module-I. Figure 1 illustrates the MMIP workflow. Users can select the appropriate module based on their experimental design and data. New users are encouraged to consult the MMIP manual, example data, and results. A job queue system notifies users via email with a job-based ID and access link for analysis results.

**Figure 1:**
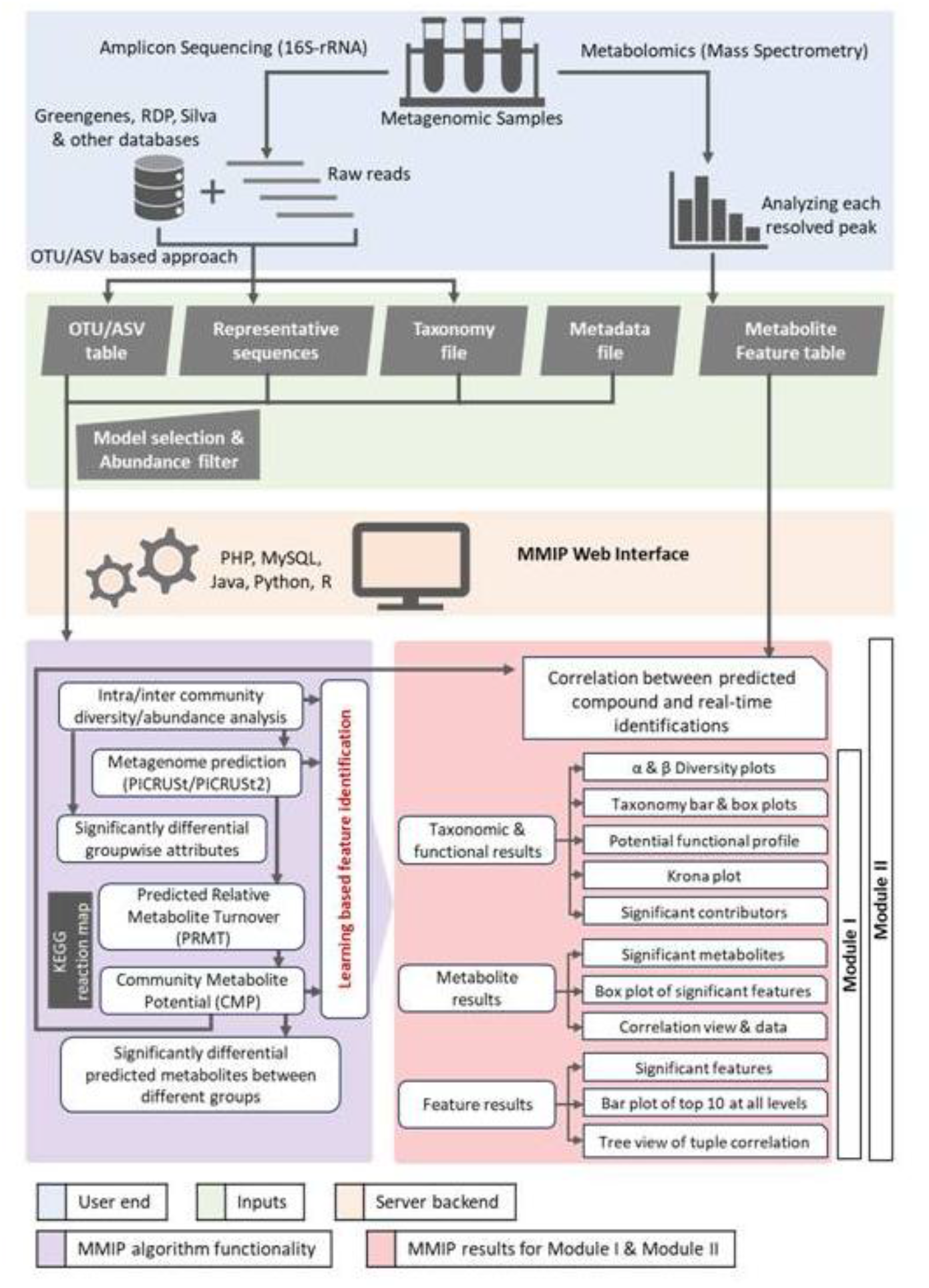
The entire workflow that runs at the backend of MMIP. Keys define different categories for each section of workflow.

### MMIP data inputs

For MMIP Modules I and II, the required file types are the OTU/ASV abundance table from 16S rRNA amplicon sequencing data and sample-wise metadata file in QIIME format (23). The abundance table can be generated by closed-reference otu-picking using Greengenes database (versions 13_5 and 13_8) or denoising with other database classifiers (such as Greengenes, SILVA, GTDB); the latter requires representative sequences and taxonomy files. Users can apply abundance-based filtering to remove low-count features. The MMIP feature selection model (Support Vector Machine, Random Forest, or Decision Tree) can be specified by the user. For Module-II, users are required to provide an extra upload of the metabolic feature table in a tab-delimited “*.txt*” file with sample names in columns and compound names in rows.

### MMIP Algorithms

#### Estimation of community-wise metabolic potential (CMP)

We predicted metagenome content from 16S rRNA amplicon data using PICRUSt (24) or PICRUSt2 (25). We normalized the OTU/ASV abundance table by predicted 16S copy number and performed metagenomic function prediction. This involved a gene content inference step, generating a table (matrix *A_nk_*, where n-rows represent the predicted genes, KEGG Orthology or KO identifiers, while k-columns represent all the samples) of predicted gene family counts for each sample. The metagenome inference step provided values representing the contribution of mapped OTUs/ASVs to their gene families. For community-wise metabolite prediction, we used the method by Noecker *et al*. (19), which requires the predicted gene list (KO identifiers) as input. The KO identifiers are mapped to the KEGG reaction map using a surjective function and based upon the directionality of the mapped reactions. Further, these reactions are mapped to the metabolic pathways. A matrix *B_mn_* of dimensions *m*×*n* was developed using the reaction stoichiometry, where m (rows) are the compound identifiers and n (columns) represents the KO identifiers (enzymes/genes) and each of the cells stores the stoichiometric information; it implies that if a compound is present at the reactant side, its stoichiometry will be placed in the respective cell of the table with a negative sign and if it lies at the product side its stoichiometry will be placed with a positive sign. If a particular compound is used by a KEGG enzyme in other pathways then its stoichiometric values will be added or subtracted in the table, depending on the position of the compound in that particular reaction (reactant/product side). The stoichiometric matrix *B_mn_* is then normalized, such that the sum total of positive vectors in a row is +1 and that of the negative vectors is −1 resulting in the matrix *C_mn_*.

The multiplication of *C_mn_* (*m*×*n*) matrix with that of the predicted metagenome function matrix *A_nk_* (*n*×*k*) results in a Community Metabolic Potential (CMP) score matrix *D_mk_*having community-wise metabolic potential for each of the metabolites/compounds in the corresponding samples, where m (rows) and k (columns) represent the metabolites and the sample names respectively. The CMP score, much like PRMT (18), is unit-less that only predicts the relative capacity of the microbial community to produce or to consume each metabolite. The mathematical expression of the CMP calculation is shown in supplementary Figure 1.

#### Feature prediction using machine learning

The CMP matrix, as mentioned previously, is formed at the backend, and is then used along with the OTU/ASV table and the predicted metagenome table, to predict features that differentiate between the sample groups. MMIP allows the user freedom to opt for any model for the feature selection – including SVM, Random Forest, and Decision Tree, which are popular models used for microbiome data (26–28). This is the most unique attribute of MMIP and is explained in Figure 2.

**Figure 2:**
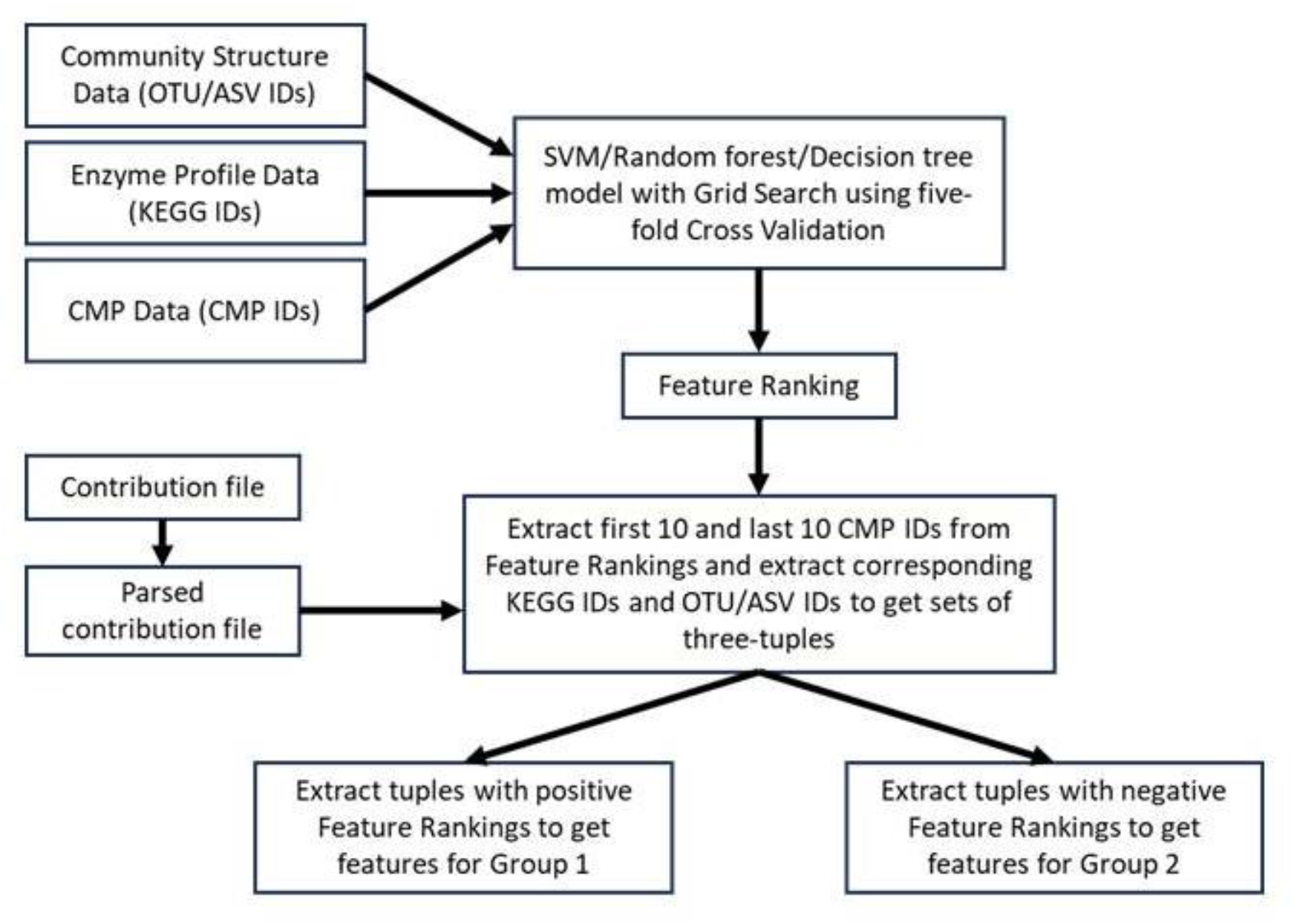
Flowchart for the classification and feature extraction of the two groups for MMIP.

A Support Vector Machine (SVM) was used to predict the hyperplane separating the data points in a multi-dimensional space. For each dataset, a gene prediction table, a compound metabolic potential score table and a metagenome abundance table (OTU/ASV table) with sample-specific data were generated and all were concatenated sample-wise in order to perform classification. The mathematical formulation of SVM can be stated according to Equation 1 with constraints as per Equation 2 where x represents the variables, y is the class outcome where y∈{-1,+1} (for an SVM model having two classes), C is a hyperparameter used for regularization, w represents the coefficient vector, b is a constant, ξ is the set of slack variables and φ represents the kernel used for data transformation.

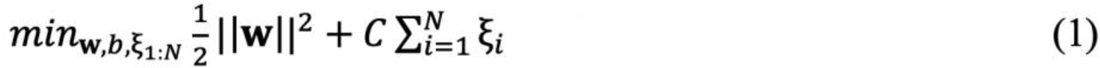

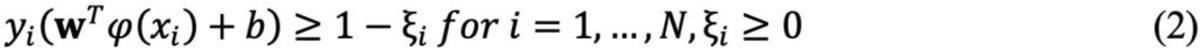

Random forest (29) is a model where a set of decision trees is used on a dataset. The model is set up so that the error converges with increase in the number of trees in the forest. Random forest models use feature selection to decide the split at every node. A decision tree (30) is the simpler version of random forest where a single tree is used with splits at every node to determine the outcome of classification based on decisions at the various nodes. For all the mentioned machine learning algorithms (31, 32), we use the Python implementation in scikit-learn. The decision tree in this implementation is of the CART (Classification and Regression Trees) algorithm (33) that seeks to maximize information gain at node *A* (*gain*(*A*); Equation 3.3), which is defined as the difference between the information *I* (*I*(*p, n*); Equation 3.1) and the expected weighted information *E*(A) (*E*(*A*); Equation 3.2) (30).

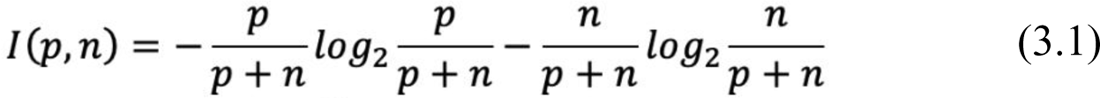

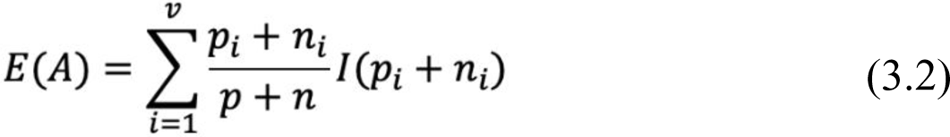

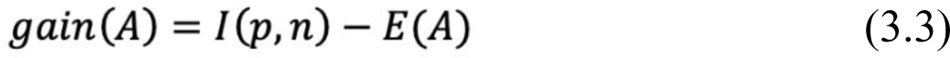

In terms of each machine-learning algorithm, a grid search using the five-fold cross-validation approach is carried out on the whole dataset to pick the hyperparameters that enable the model to have the best performance among a number of combinations of hyperparameters, implemented mainly by sklearn Python package. More specifically, for SVM, we check the values of gamma=(1e-3,1e-4), and C=(1,10,100,1000) with linear kernel and probability=True options. For random forest, we check n_estimators=(10,50,100) and criterion=(gini, entropy). For the decision tree, we also check criterion (gini, entropy). The model configured with the determined combination of hyperparameters is further fitted on the whole dataset, in order to track the specific features that have a significant contribution to each category of the target dataset.

A nested cross-validation approach with both inner folds and outer folds of 5 was performed on the whole dataset to provide a robust generalization performance evaluation for each machine learning model, since its architecture have the advantage of preventing the model from overfitting on the dataset with limited size. The evaluation metrics including AUC, accuracy, recall, F1-score, and precision which are calculated on the merged output from each outer fold are used to reflect a single model’s ability on classifying the target dataset.

### MMIP analyses

#### Functional profiling and metabolite potential

A Python pipeline performs sanity checks (removal of singleton OTUs/ASVs, normalization by rRNA copy number etc) and comparative metagenome prediction (PICRUSt/PICRUSt2), calculating CMP scores. An R function conducts differential analysis at KEGG functional and CMP levels using statistical tests such as Wilcoxon test, ANOVA, t-test etc. Predicted metagenome results, CMP compound-wise boxplots, and statistical analysis are accessible to the user with proper visualization and downloadable diagrams using Plotly (34), D3 (35), Krona Plot (36) and flat file on the web interface

#### Community profiling and diversity analyses

The user-uploaded OTU/ASV table and metadata file is processed through a R pipeline, which also uses the Phyloseq package (37) for comparative taxonomic profiling, alpha diversity analyses (Shannon, Simpson diversity indices, effective numbers) and beta diversity analyses (Bray-Curtis dissimilarity, GUniFrac (38), UniFrac-weighted/unweighted). Further the R function is supposed to perform the differential analysis at different taxonomic levels along with all the statistical analysis including Wilcoxon test, Kruskal-Wallis, ANOVA, t-test etc. All results are available for the user visualization and download.

#### Feature selection and predictive linkage identification

The top ten positive and negative CMP features were used along with contribution file in order to link them with corresponding KEGG enzymes and OTUs/ASVs. Through multiple iterations a threshold of 0.7 was selected for the presence of OTUs/ASVs in a certain number of samples. This implies that the corresponding mean OTU/ASV contributed would reflect a certain quantity only if the number of samples for that OTU/ASV exceeded the threshold value, otherwise the value in the mean columns will be zero. The features are then linked according to their feature ranking obtained earlier. The three-tuples that have positive ranks for all three features (the linked CMP, KEGG enzyme and OTU/ASV) have a greater probability of contributing to the positive side of the hyperplane and hence represent collective features for this particular group. Likewise, it can be said for the negatively ranked three-tuples that represent collective features for the other group considered for classification.

#### Correlation of predicted vs real time metabolites

This functionality is only available with Module-II. Here, users can easily correlate predicted metabolic information with real-time metabolite data (LC-MS, GC-MS feature table). The framework identifies common compounds between predicted and real-time data, applying the Mantel test on their distance matrices. It determines whether the predicted and matched data are positively or negatively correlated based on the correlation coefficient (−1 to +1 range). This allows users to identify the most probable microbial source for the compounds or metabolites detected in the real-time metabolite data.

#### Dataset selection for validation of our webtool

In order to validate our platform, we selected three studies with publicly available amplicon sequencing data coupled with metabolome data. These validation datasets are from the studies of Srinivasan *et al.* (2015) (dataset-I), Dhakan *et al*. (2019) (dataset-II), and Jacobs *et al*. (2023) (dataset-III) (39–41). Amplicon sequencing data (accession numbers PRJNA275907, PRJNA397112, PRJNA812699) and paired mass spectrometry data were obtained. All the amplicon data was re-analysed using standard QIIME/QIIME2 pipeline (23, 42), and the OTU/ASV table was obtained either by closed-reference OTU picking against Greengenes v13.8 database (43, 44) or denoised with DADA2 q2 plugin and then assigned taxonomy using SILVA v138 classifier (45). The OTU/ASV table and associated metadata in QIIME format were uploaded to MMIP for analysis. MMIP predicted metabolites were compared with reported data to assess concordance with the prediction model. We used Module-I for dataset–I and Module-II for dataset-II and dataset-III. Metabolomics data were processed and annotated with KEGG IDs separately for dataset II and III (Additional File 1).

## Result

### Comparative community-wise profile analyses for validation dataset-I

The cohort B validation dataset-I (Srinivasan *et al.* (2015) study) contains 40 Bacterial Vaginosis (BV) samples and 20 healthy (H) samples. The alpha diversity (Shannon) analysis by MMIP accurately reflects significantly increased microbial diversity in BV samples compared to the H (Supplementary Figure 2a). Further, the beta diversity of the samples, shown in supplementary Figure 2b, depicts statistically significant separation between the H and the BV samples (p<0.001).

The taxonomic distribution showed concordance in the findings generated from MMIP, our standardised analysis and the study itself (supplementary Figure 3). It is known that *Lactobacillus* belonging to the Phylum Firmicutes, is the most abundant microbial genus found in a healthy vagina (46, 47) and it maintains the acidic environment by the production of lactic acid or lactate. MMIP results show a similar trend as reported in the study where the genus *Lactobacillus* and Phylum Firmicutes is over-abundant in the H group, while a higher percentage of diverse bacteria (*Megasphaera*, *Gardnerella*, *Prevotella* and *Sneathia*, and also *Atopobium vaginae*) were present in BV samples, indicating a dysbiotic vaginal environment.

### Comparative pathway and predicted metabolite analysis for validation dataset-I

Srinivasan *et al*. (2015) reported 108 metabolites with KEGG annotations; MMIP predicted metabolites overlapped with 52 of the reported metabolites (Supplementary Figure 10a, Supplementary Table 1). In case of metabolites prediction, MMIP provides a CMP score corresponding to each metabolite, which signifies the capacity of a bacterial community to consume or accumulate that metabolite. It showed that the relative capacity of a particular metabolite, cadaverine, being accumulated in the BV samples was significantly higher than in the healthy samples, and which agrees with the findings of the study.

The study revealed a number of metabolites including amino acids, dipeptides, and metabolites associated with carbohydrate metabolism. They reported 19 amino acids, out of which 18 were found to be in lower amounts in BV samples as compared to healthy. MMIP also predicted decreased levels of alanine, cysteine, histidine, tryptophan, phenylalanine and the dipeptide, alanylalanine, in BV samples. For metabolites related to carbohydrate metabolism the study demonstrated lowering of simple sugars, intermediate metabolite like glucose-6-phosphate, and sugar alcohols like sorbitol and mannitol in BV samples with respect to healthy. MMIP not only was able to predict all the above-mentioned compounds but also highlighted their tendency to be lower in the BV samples. In addition to that, the study reported lowering of glycerol, ethanolamine and glycerophosphorylcholine (GPC), all of which are associated with lipid metabolism, in BV samples and they too were correctly predicted by MMIP.

### Feature link prediction for validation dataset-I

MMIP predicts links between OTUs, KEGG IDs (enzyme profiles), and metabolites. It is noteworthy that CMP scores in MMIP does not indicate net usage or synthesis of metabolites; positive score suggests increased utilization and negative score implies more accumulation when comparing sample groups.

Such positive and negative feature links obtained from MMIP on -validation dataset-I is depicted in supplementary Figure 4a and 4b respectively. The positive link is predicted for the compound C00089 (sucrose), where it is associated with the enzyme, sucrose phosphorylase (K00690) and further downstream to *Lactobacillus* species. *Lactobacillus* is abundant in the healthy (H) samples, it can thus be expected that a high *Lactobacillus* population causes an increased utilization of the metabolite via the sucrose phosphorylase enzyme. The negative link is predicted for the compound C00436 (N-Carbamoylputrescine), which is associated with the enzyme agmatine deiminase (K10536) and further with the bacterial family *Coriobacteriaceae*, harbouring genera like *Eggerthella* sp. and *Atopobium* sp., specific to the BV samples according to the study. The MMIP link thus suggests that increase in these particular bacterial species in BV group would mean increased action of the enzyme agmatine deiminase, yielding a higher amount of the metabolite Carbamoylputrescine in the samples.

### Comparative community-wise profile analyses for validation dataset-II

Validation dataset-II (Dhakan *et al*. (2019) study) comprised 110 individuals with omnivorous and vegetarian diet, belonging to the Southern and the North-Central states of India, respectively. We filtered out samples with less than 1% of maximum OTU, resulting in a final dataset of 57 omnivorous and 35 vegetarian samples and uploaded to MMIP. The genus-level taxonomic plot generated by MMIP demonstrates a clear difference between the two groups (Supplementary Figure 5). The significant OTUs obtained from MMIP for this dataset correlated with the findings of the original study. The results showed that the omnivorous group was enriched in *Ruminococcus* and *Faecalibacterium* (including *F. prausnitzii*), and *Roseburia* and *Akkermensia muciniphila*; while the vegetarian group had bacterial genera *Prevotella* and *Megasphaera*.

### Comparative pathway and predicted metabolite analysis for validation dataset-II

Dhakan *et al*. (2019) highlighted enrichment of 5928 KEGG pathways forming two clusters associated with two different diet-groups. MMIP also generated KEGG pathway level feature prediction distinguishing between the two groups and 4918 of those overlapped with that of the study. The study also identified 13 annotated metabolite clusters, 6 of which overlapped with MMIP results (Supplementary Table 2). We processed the metabolomic data separately, identify 306 metabolites, out of which 103 could be annotated and 53 matched with our metabolite prediction (Supplementary Figure 10b, Supplementary Table 3).

While analysing the individual trends of the predicted metabolites, MMIP predicted many of the metabolites as mentioned in the study, including serine (C00065), butyrate (C00246) and valine (C00183), all of which were found to be associated with that of the omnivorous diet group. For the metabolite serine, MMIP assigned a high negative CMP score in the omnivorous group, implying that the omnivorous group associated microbes have a relatively higher potential of serine synthesis; the same holds true for valine as well. SFCAs (short chain fatty acids) like butyrate, propionate, correlate with that of omnivorous diet, and butyrate is found to have a negligible CMP score in vegetarian group, while many of the samples of the omnivorous group show a negative CMP score concluding that again this group has a higher potential of synthesizing butyrate. All the above-mentioned results from MMIP follow the same trend as reported in the study.

### Feature link prediction for validation dataset-II

MMIP predicted dietary habit specific links for validation dataset-II. The positive link is predicted for the compound manninotriose (C05404), and then linked with the enzyme alpha-galactosidase (K07406) and the members of *Ruminococcaceae* family and of genus *Ruminococcus* (Supplementary Figure 6a). These microbes are related to the omnivorous diet group and could imply the increased utilization of the associated metabolite. Similarly, another link was predicted for the compound sucrose-6-phosphate (C02591) associated with the enzyme sucrose-6F-phosphate phosphohydrolase (K07024) and in turn with microbial genera *Ruminococcus* and *Faecalibacterium*, found in omnivorous diet group, shown in supplementary Figure 6b. This linkage too shows increased utilization of the metabolite via the enzyme in the given set of microbes.

### Comparative community-wise profile analyses for validation dataset-III

Validation dataset-III, Jacobs *et al*. (2023) investigated patients with Irritable Bowel Syndrome (IBS) and healthy controls. We specifically selected the cross-sectional samples that included paired metabolomic data without repeats for our analysis. Our final dataset consisted of 181 and 134 samples from individuals with IBS and control subjects respectively. MMIP analysis with ASV table yielded non-significant differences in alpha diversity, similar to the findings in the study, although, differences in beta diversity were significant and the R^2^ values were comparable, as also found in the study (Supplementary Figure 7).

The significant taxa identified in MMIP were consistent with the findings of the study. In the IBS group, there was a significant abundance of *Actinomyces*, *Bacteroides*, *Alistipes*, *Faecalibacterium*, and *Blautia hydrogenotrophica* species. Furthermore, the family *Oscillospiraceae* and *Ruminococcaceae*, along with the class Clostridia, exhibited increased population. In contrast, the genera *Bilophila*, *Roseburia*, and *Subdoligranulum* were significantly reduced in the IBS group. Refer to supplementary Figure 8 for a taxonomic plot comparing the genera between the two groups.

### Comparative predicted metabolite analysis for validation dataset-III

Jacobs *et al*. (2023) conducted an extensive analysis of metabolite profiles, and we successfully matched KEGG compound IDs to 306 of those metabolites. By comparing MMIP predicted metabolites with those matched ones, we found 134 overlapping compounds (Supplementary Figure 10c, Supplementary Table 4). Especially, tyramine, which is known for its association with promoting IBS, and fumarate, were found to be elevated in IBS samples. Conversely, hydrocinnamate and 2-oxoarginine were observed in lower quantities compared to control samples.

### Feature link prediction for validation dataset-III

MMIP predicted connections between metabolites, enzymes, and microbial community features that were congruent with IBS-related characteristics found in the study. The first link involved Hematoside (G00108), Hexosaminidase (K12373) enzyme, and *Actinomyces sp*., *Alistipes sp*., *Bacteroides fluxus*, and *Prevotella timonensis* (Supplementary Figure 9a). The study reported an abundance of these bacteria, elevated levels of the enzyme, and increased glycan metabolism in IBS samples. The reaction shown in this link also is part of the glycan metabolism catalyzed by the Hexosaminidase enzyme. The second link connected Lactosylceramide (C01290), Beta-galactosidase (K01190) enzyme, and *Actinomyces* sp., *Faecalibacterium* sp., *Bacteroides* sp., and *Blautia hydrogenotrophica* (Supplementary Figure 9b). The feature compound in the first link had higher accumulation in the IBS group, while the second link’s feature compound had lower accumulation. These linked bacterial groups, as well as the Beta-galactosidase enzyme, were specific to IBS, suggesting increased forward reaction in the IBS samples.

### Run time, model performance and comparison with other online tools

The runtime of MMIP’s Module-I depends on the biom file size and number of samples. Feature density in the biom file also affects the processing time. We selected two datasets, Srinivasn *et al*. (2015) and Armstrong *et al*. (2018) (48), created subsets, processed reads with QIIME1 and QIIME2, and uploaded the data individually. Supplementary Table 5 provides the time requirements for different file types, and shows that time increases linearly with sample count.

For feature prediction in the validation datasets, SVM algorithm was used as it excels in high-dimensional spaces. Evaluation of model performance shows that SVM and RF perform similarly, with SVM outperforming for the third dataset. Further we have included metrics for all the models (Supplementary Table 6), ROC curves (Supplementary Figure 11), and confusion matrices (Supplementary Figure 12) to showcase overall performance.

We performed a qualitative comparison of MMIP with few other online microbiome analysis tools - EasyMAP (49), M2IA (50), MicrobiomeAnalyst 2.0 (51, 52) and VAMPS (53), highlighting their different features and functionalities (Supplementary Table 7). While these tools offer comprehensive microbiome and statistical analysis, they lack knowledge-based integration of microbiome and metabolome data. We also performed benchmarking of MMIP with MicrobiomeAnalyst 2.0 using all three datasets (Supplementary Table 8).

## Discussion

We validated the functionality of MMIP using datasets from three different studies. Using data from Srinivasan *et al* (2015) (39), MMIP predicted 52 metabolites that overlapped with real time metabolites from the study, along with microbial profiling and diversity indices comparison. Supplementary Figure 4a explains how the positive link could be interpreted biologically and biochemically. Sucrose (C00089) is a disaccharide made of fructose and glucose and the enzyme sucrose phosphorylase (K00690, part of carbohydrate metabolism reaction map), breaks it down to form fructose and glucose-1-phosphate (KEGG reaction ID R00803). This enzyme is further associated with *Lactobacillus iners* and other *Lactobacillus* species, according to MMIP. As per the study, these microbes are associated with the healthy group. Increase in the utilization of sucrose (C00089) is shown by the high positive CMP score in healthy group, suggesting that the forward reaction is taking place and hence there would be an increased formation of fructose, which was found to be less in BV group in the study. Similarly, the reaction in the supplementary Figure 4b suggests, the increase in the population of BV associated bacteria might be associated with an increase in the enzyme action of agmatine deiminase (K10536), along with the entire Arginine degradation pathway, allowing the forward reaction and hence more production of N-Carbamoylputrescine (C00436), a precursor for putrescine, which is a compound elevated in BV samples in the study. The CMP score of N-Carbamoylputrescine shows that this compound is highly accumulated in BV. It can thus be concluded that this reaction is possibly quite important for BV associated bacteria.

In our second validation dataset by Dhakan *et al*. (2019) (40), the taxonomic distribution between the two groups by MMIP was at par with that of the study, showing significant presence of bacteria *Roseburia* and *Akkermensia muciniphila* in the omnivorous group and that of *Prevotella* and *Megasphaera* in the vegetarian group. The study also checked pathways associated with the two diet groups and when compared with MMIP results they showed remarkable overlap with our pathway analysis. MMIP also predicted the feature linkages, one of them describing the increase in the omnivorous diet specific microbes, which is related to higher activity of the alpha-galactosidase enzyme (K07406), thus favouring forward reaction that depletes manninotriose while increasing D-Galactose (C00124) in the system (Supplementary Figure 6a). Another feature points out that omnivorous group is enriched in Phylum Firmicutes, genus *Ruminococcus*, and species *Faecalibacterium prausnitzii*, implying an increased action of the associated enzymes. Thus, an elevated conversion from sucrose–6-phosphate (C02591) to sucrose (C00089) and eventually to D-Glucose (C00031) and D-Fructose (C00095), both of which were reported to be increased in omnivorous diet (Supplementary Figure 6b).

The final validation study was by Jacob *et al*. (2023) (41) and MMIP was able to reproduce the microbial population signatures mentioned in the study, including *Actinomyces*, *Bacteroides*, *Alistipes*, *Faecalibacterium* and *Blautia hydrogenotrophica* being significantly abundant in IBS samples while lowering of *Bilophila*, *Roseburi*a and *Subdoligranulum*. With respect to metabolites, MMIP was able to predict many of the IBS associated metabolites including tyramine which increased in IBS and hydrocinnamate and 2-oxoarginine, which were lowered. Finally, the MMIP predicted links that highlighted two compounds, one of which accumulated in IBS samples while the other depleted. The first link shows that the bacteria associated with IBS (like *Bacteroides fluxus*, *Prevotella timonensis*) were linked with the Hexosaminidase enzyme (K12373) which drives the forward reaction changing Ganglioside (G00109) to Hematoside (G00108), causing an accumulation of the latter in IBS samples, as indicated by the lower CMP score (Supplementary Figure 9a). The fact that the IBS associated bacteria were reported to have increased Glycan metabolism and the reaction mentioned in the link (KEGG reaction ID R06004), is part of the Glycan metabolism pathway, proves the validity of the link. The second link associates another set of IBS bacteria, *Actinomyces*, *Bacteroides* and *Blautia hydrogenotrophica*, with Beta-galactosidase enzyme, which drives the reaction (KEGG reaction ID R03355) forward, depleting the compound Lactosylceramide (C01290), and which is also shown by the higher negative CMP score for IBS (Supplementary Figure 9b).

When using MMIP, it is important to consider key points and limitations to ensure proper interpretation of the results. MMIP provides the potential value of metabolite utilization within the microbial community, rather than absolute content or metabolism rate. The predicted metabolites in MMIP require experimental validation. Data pre-processing using standardized pipelines is necessary for reliable results. Limitations and biases associated with the KEGG reference database should be considered. Predicted features and linkages may lack statistical significance and should be interpreted cautiously. Users should consider the sample size, as statistical tests do not work for very small sample numbers. It is important to understand that the feature links identified by MMIP can help to generate preliminary hypotheses about the types of reactions occurring in a microbial community. However, further experimental validation and rigorous investigation are necessary to confirm these hypotheses.

This web-resource offers diversity analysis, taxonomy profiling, metagenome prediction, and metabolite potential measurement and a supervised learning-based approach to connect taxonomy abundance, enzyme profiles, and metabolite potential. It provides the user an initial idea of the probable metabolite signature from targeted amplicon sequencing data without the need for metabolomic analysis. MMIP could also help the user to form a preliminary idea of the metabolic potential or the enriched pathways of the microbial communities; and facilitates the formation of biologically valid hypotheses for comparing sample groups.

## Data and code availability

The web-server is available at https://github.com/websandip/mmip and the data for validation was from already published studies which can be accessed by NCBI BioProject through Project ID mentioned in the main text. Further detailed manual on how to use MMIP is available at https://github.com/websandip/mmip/blob/main/manual.pdf

## Supplementary data

All supplementary data (including files, figures and tables) will be available in the published version of this paper.

## Acknowledgment

The research was supported by the Ramanujan Fellowship Grant of the Science and Engineering Research Board (SERB), Government of India, awarded to SP. SP also acknowledges the support of the Bioinformatics Centre, JISIASR. DB received fellowship support from the Indian Council of Medical Research, Government of India. AG acknowledge support by the High Performance and Cloud Computing Group at the Zentrum für Datenverarbeitung of the University of Tübingen, the state of Baden-Württemberg through bwHPC, the German Research Foundation (DFG) through grant no. INST 37/935-1 FUGG. AG and WZ also acknowledge support of the BMBF-funded de.NBI Cloud within the German Network for Bioinformatics Infrastructure (de.NBI) (031A532B, 031A533A, 031A533B, 031A534A, 031A535A, 031A537A, 031A537B, 031A537C, 031A537D, 031A538A). We would like to express our gratitude to Prof. Dr. Boris Maček, Dr. Daniel Petras, and Dr. Timo Sachsenberg for their valuable discussions on GC-MS data analysis. Additionally, we extend our thanks to Prof. Dr. Nico Pfeifer and Jonas Ditz for their insights on machine learning methods.

