## Supplementary Table for "Microbiome Metabolome Integration Platform (MMIP): a web-based platform for microbiome and metabolome data integration and feature identification" (from another version of this preprint: version 1, 2023-04-06)

| **Metabolite group mentioned by the study** | **MMIP predicted metabolites that fall under the designated cluster and also following a similar trend** |
|---|---|
| MES_9_Glycerol | Glycerol |
| MES_48_L_valine_Leucine_Isoleucine_Serine | L-Serine, L-Valine |
| MES_35_Propanoic_Acid | Propanoic Acid |
| MES_19_Cadaverine_Lauric_acid | Cadaverine |

**Supplementary Table 1:** MMIP predicted metabolites that overlapped and followed the trend with respect to validation dataset 2.
