## Supplementary Figure for "Microbiome Metabolome Integration Platform (MMIP): a web-based platform for microbiome and metabolome data integration and feature identification" (from another version of this preprint: version 1, 2023-04-06)

We denote Sample-wise Gene Abundance as:

$$A_{nk} = \begin{pmatrix} a_{1,1} & a_{1,2} & \cdots & a_{1,k} \\ a_{2,1} & a_{2,2} & \cdots & a_{2,k} \\ \vdots & \vdots & \ddots & \vdots \\ a_{n,1} & a_{n,2} & \cdots & a_{n,k} \end{pmatrix}_{n \in Genes, k \in Samples} \tag{1}$$

We denote connectivity matrix of network in graph B as:

$$B_{mn} = \begin{pmatrix} b_{1,1} & b_{1,2} & \cdots & b_{1,n} \\ b_{2,1} & b_{2,2} & \cdots & b_{2,n} \\ \vdots & \vdots & \ddots & \vdots \\ b_{m,1} & b_{m,2} & \cdots & b_{m,n} \end{pmatrix}_{m \in Compound, n \in Genes} \tag{2}$$

The normalization matrix of $B_{mn}$ as shown in below:

$$C_{mn} = \begin{pmatrix} c_{1,1} & c_{1,2} & \cdots & c_{1,n} \\ c_{2,1} & c_{2,2} & \cdots & c_{2,n} \\ \vdots & \vdots & \ddots & \vdots \\ c_{m,1} & c_{m,2} & \cdots & c_{m,n} \end{pmatrix}_{m \in Compound, n \in Genes} \tag{3}$$

where

$$c_{m,n} = \begin{cases} \frac{b_{m,j}}{\sum B_m{}^+}, & b_{m,j} > 0, |B_m{}^+| > 1, j \in (1,n) \\ 1, & b_{m,j} > 0, |B_m{}^+| = 1, j \in (1,n) \\ 0, & b_{m,n} = 0 \\ \frac{b_{m,j}}{|\sum B_m{}^-|}, & b_{m,j} < 0, |B_m{}^-| > 1, j \in (1,n) \\ -1, & b_{m,j} < 0, |B_m{}^-| = 1, j \in (1,n) \end{cases}$$

and $B_m{}^+ = \{b_{mj} \mid b_{mj} > 0\}$, $B_m{}^- = \{b_{mj} \mid b_{mj} < 0\}$.

CMP Score obtained by multiplying Environmental Metabolome Matrix with Sample-wise Gene Abundance. This process is described as below:

$$\begin{aligned} D_{mk} &= C_{mn} \times A_{nk} \\ &= \begin{pmatrix} c_{1,1} & c_{1,2} & \cdots & c_{1,n} \\ c_{2,1} & c_{2,2} & \cdots & c_{2,n} \\ \vdots & \vdots & \ddots & \vdots \\ c_{m,1} & c_{m,2} & \cdots & c_{m,n} \end{pmatrix} \times \begin{pmatrix} a_{1,1} & a_{1,2} & \cdots & a_{1,k} \\ a_{2,1} & a_{2,2} & \cdots & a_{2,k} \\ \vdots & \vdots & \ddots & \vdots \\ a_{n,1} & a_{n,2} & \cdots & a_{n,k} \end{pmatrix} \\ &= \begin{pmatrix} d_{1,1} & d_{1,2} & \cdots & d_{1,k} \\ d_{2,1} & d_{2,2} & \cdots & d_{2,k} \\ \vdots & \vdots & \ddots & \vdots \\ d_{m,1} & d_{m,2} & \cdots & d_{m,k} \end{pmatrix}_{m \in Compound, k \in Samples} \end{aligned} \tag{4}$$

Supplementary Figure 1: Calculation of the algorithm for CMP score generation by Noecker et al which works in the backend of the server.

a

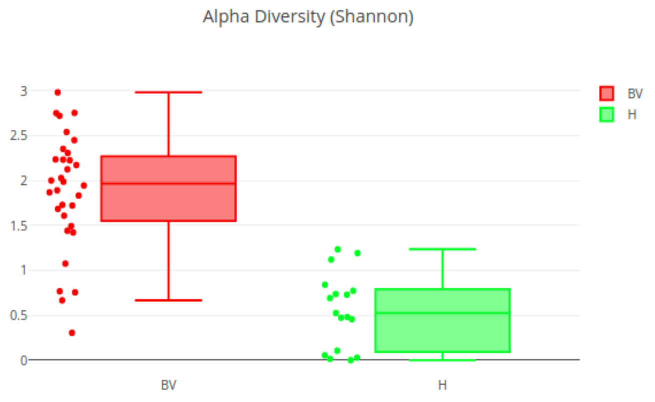


| Diversity Indices | Significance Test | MMIP | Separately calculated |
|---|---|---|---|
| Shannon | ANOVA | 9.93e-10 | 9.3e-12 |
| Simpson | ANOVA | 1.03e-08 | 3.31e-07 |

b

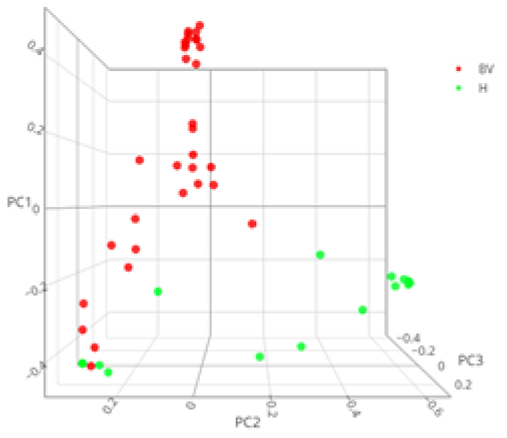


| Adonis Test | MMIP | Separately calculated |
|---|---|---|
| F-statistics | 13.433 | 13.541 |
| P-value | 0.001 | 0.001 |

Supplementary Figure 2: Here we show (a) Alpha diversity comparison along with the different indices performed by MMIP and our own separate analysis and (b) the beta diversity among healthy and disease group performed by MMIP.

a

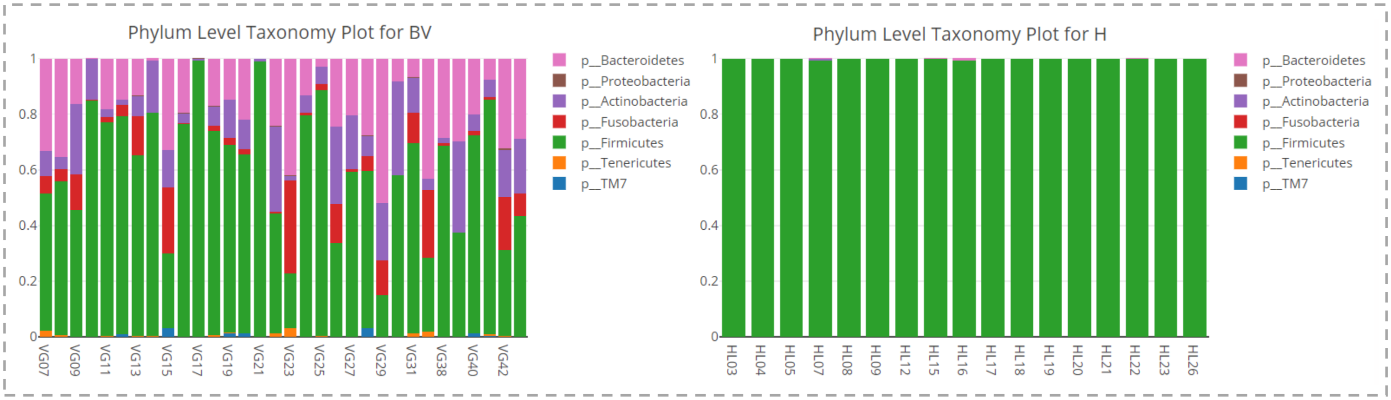


b

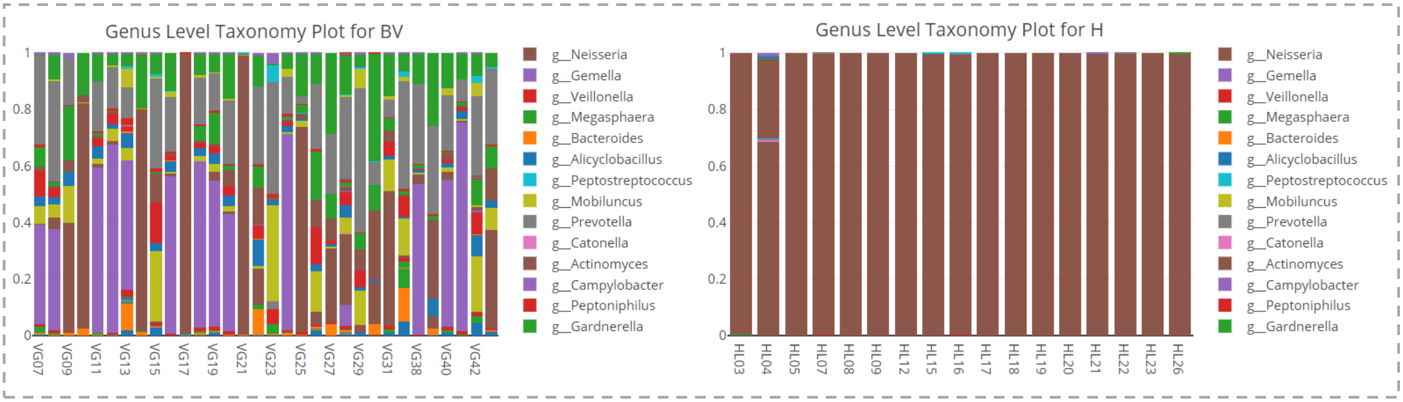


Supplementary Figure 3: Taxonomic profile showing both phylum-wise (a) and genus-wise (b) separation between BV and Healthy (H) groups performed by MMIP.

a

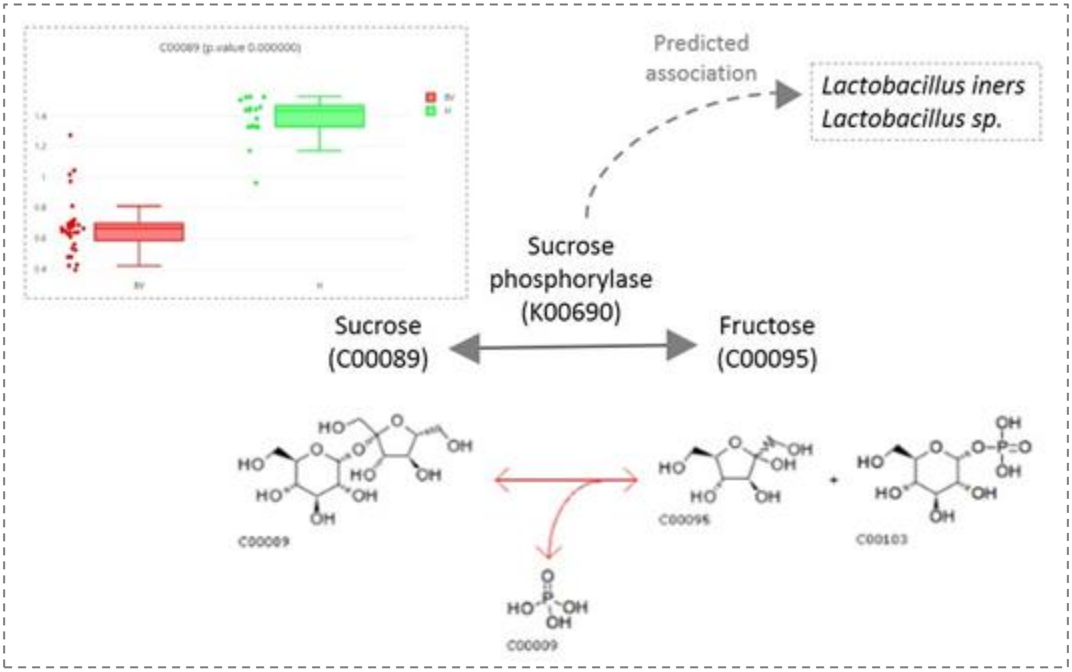


b

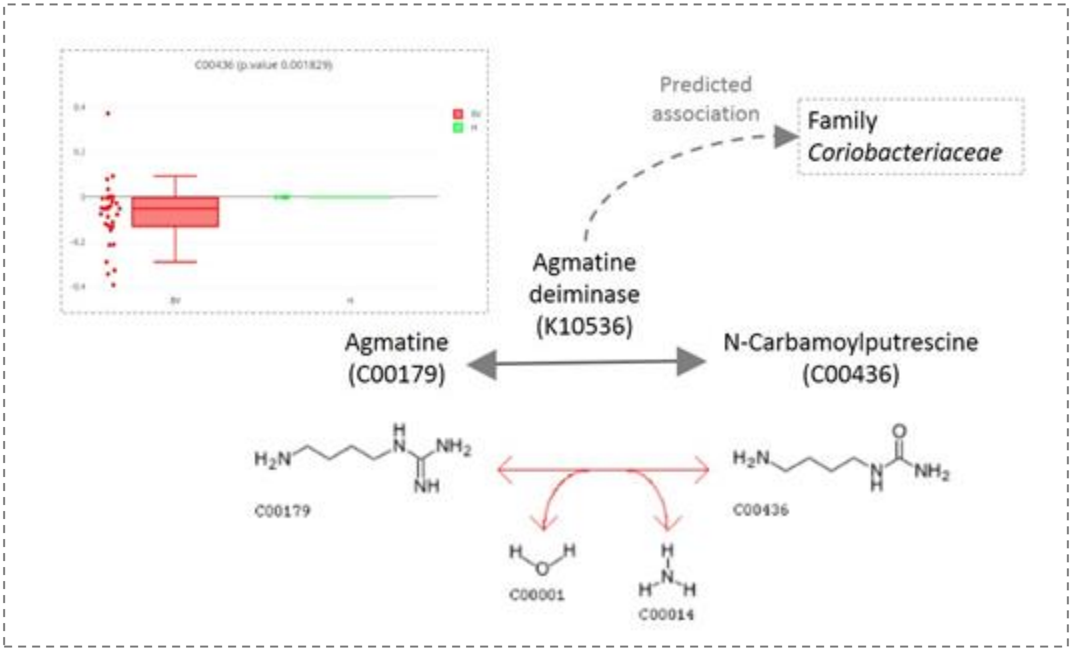


Supplementary Figure 4: Positive (a) and negative (b) linkages predicted by MMIP on validation dataset 1.

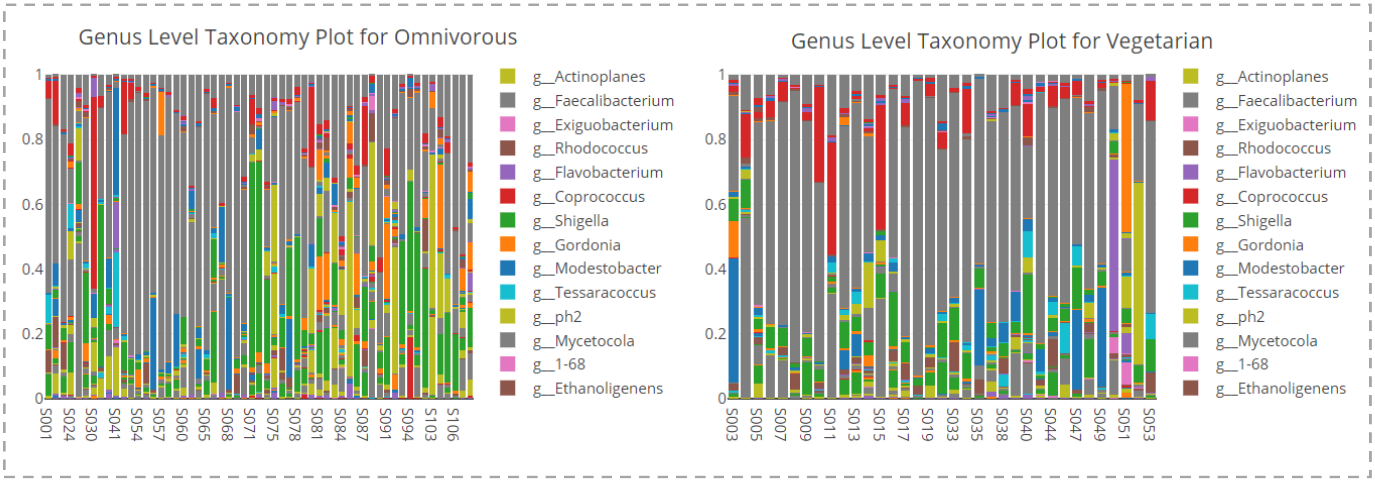


Supplementary Figure 5: Taxonomic profile showing genus-wise separation between omnivorous and vegetarian groups preformed by MMIP.

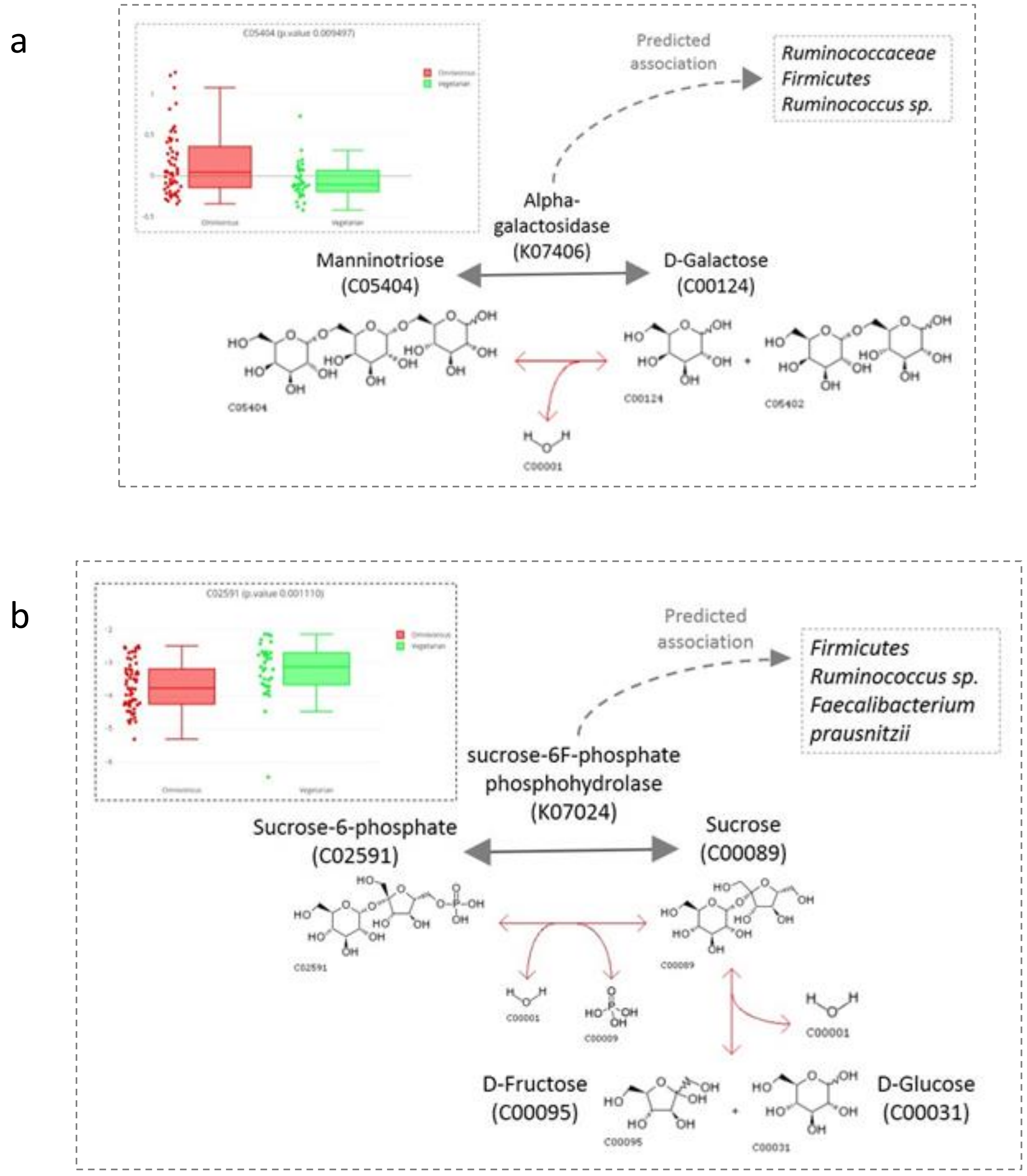


Supplementary Figure 6: Positive (a) and negative (b) linkages predicted by MMIP on validation dataset 2.
